# AllerTrans: A Deep Learning Method for Predicting the Allergenicity of Protein Sequences

**DOI:** 10.1101/2024.08.09.607419

**Authors:** Faezeh Sarlakifar, Hamed Malek, Najaf Allahyari Fard, Zahra Khotanlou

## Abstract

Recognizing the potential allergenicity of proteins is essential for ensuring their safety. Allergens are a major concern in determining protein safety, especially with the increasing use of recombinant proteins in new medical products. These proteins need careful allergenicity assessment to guarantee their safety. However, traditional laboratory testing for allergenicity is expensive and time-consuming. To address this challenge, bioinformatics offers efficient and cost-effective alternatives for predicting protein allergenicity. In this study, we developed an enhanced deep-learning model to predict the potential allergenicity of proteins based on their primary structure represented as protein sequences. In simple terms, this model classifies proteins into allergenic or non-allergenic classes. Our approach utilizes two protein language models to extract distinct feature vectors for each sequence, which are then input into a deep neural network model for classification. Each feature vector represents a specific aspect of the protein sequence, and combining them enhances the outcomes. Finally, we effectively combined the predictions of our top-performing models using ensemble modeling techniques. This could balance the model’s sensitivity and specificity and improve the outcome. Our proposed model demonstrates admissible improvement compared to existing models, achieving a sensitivity of 97.91%, specificity of 97.69%, accuracy of 97.80%, and an impressive area under the ROC curve of 99% using the standard five-fold cross-validation.

## 1. Introduction

Proteins play an important role in vital cellular processes. Any disruption in protein function can lead to disease. Since the advent of medical sciences, there has been a continuous effort to find and develop effective treatments. In recent decades, with the advancement of technology, novel methods have emerged to help researchers produce pharmaceuticals by generating recombinant proteins. The application of recombinant proteins to treat major diseases like hepatitis, cancer, and diabetes has greatly revolutionized modern medical science. Due to the advantages of recombinant proteins over chemical medicines, the global demand for them is rising. Thus, the annual sale of such medical drugs has been over $2 billion. Projections suggest this market will grow to around $9 billion by 2032 [1]. In addition, recombinant proteins are widely used in industry and agriculture.

Studying the safety of proteins, especially recombinant proteins, is essential for their effective utilization. One of the most important safety considerations is assessing allergenicity and ensuring proteins are non-allergen. Allergies and allergens affect about one-third of the world’s population. According to a report by the World Allergy Organization, about 10 to 30 percent of the worldwide adult population is affected by allergic rhinitis. Additionally, adverse drug reactions may affect up to 10% of the world’s population and up to 20% of all hospitalized patients, with drugs potentially responsible for up to 20% of fatalities due to anaphylaxis. Furthermore, sensitization rates to one or more common allergens among school children are approaching 40%-50% [2]. Therefore, investigating and identifying whether proteins are allergenic or non-allergenic is greatly important.

Bioinformatics is an interdisciplinary concept that has emerged from the combination of biological and computer sciences and is used in many biological research. Generally, two methods of laboratory testing and bioinformatics analysis are used to identify the allergenicity of proteins. Since the laboratory testing process is time-consuming and expensive, bioinformatics analyses have received huge attention in this area.

Bioinformatics methods for allergenicity prediction of protein sequence can be categorized into three groups: traditional similarity-based methods, statistical methods based on biological protein features, and deep-learning-based methods. In bioinformatics studies for protein allergenicity prediction, researchers often utilize one or a combination of these three methods.

### Similarity-Based Methods

These methods traditionally involve assessing protein sequence similarity using tools such as the Basic Local Alignment Search Tool (BLAST) [3]. They compare protein sequences against local or online databases to identify similarities with known allergens. However, their efficiency can be limited by the requirement for similar allergenic protein counterparts and reduced performance with large-scale data.

### Statistical Methods Based on Biological Protein Features

These methods involve extracting biological features from protein sequences, such as amino acid composition, and encoding them into numerical representations using statistical techniques. These features are subsequently employed with prediction methods, such as machine-learning models, to predict the allergenicity of protein sequences. Unlike traditional search-based methods that primarily rely on sequence similarity, these approaches leverage statistical methods to interpret biological data effectively, offering a more comprehensive approach.

### Deep Learning-Based Methods

Deep learning-based methods for protein allergenicity prediction leverage artificial neural networks to automatically learn intricate patterns and representations from protein sequences. These methods are proficient in extracting high-level features that may not be discernible through traditional or statistical approaches alone. Each of the architectures—Convolutional Neural Networks (CNN), Recurrent neural networks (RNN), and transformers—offers unique strengths in protein allergenicity prediction. CNNs are adept at capturing local patterns, RNNs excel in modeling sequential dependencies, and transformers leverage self-attention for global context modeling. While transformers are primarily used in natural language processing (NLP), their ability to represent sequences of letters makes them adaptable for protein sequence analysis as well. Transformers, in particular, have shown impressive results in this area by improving feature extraction. This improvement has boosted the model’s ability to detect subtle differences and complex relationships in protein sequences, resulting in better generalization.

By leveraging deep-learning techniques, we anticipated achieving admissible accuracy in classifying protein sequences into allergenic and non-allergenic categories, and indeed, this approach has proven to be effective, offering a promising alternative to traditional methods [4].

### 1.1 Related Work

#### 1.1.1 AllerDET

In this research study, the authors used a similarity-based method called Pairwise Sequence Alignment (PSA) for feature extraction and identified four key features. These features were then used to classify proteins into allergenic and non-allergenic categories. A combination of Restricted Boltzmann Machines and Decision Trees was used for the classification [5].

#### 1.1.2 AllerCatPro 2.0

In this work, the authors predict the 3D structure of a protein from its primary structure (protein sequence) and check if the predicted 3D structure of the input protein sequence is similar to the known 3D structures of allergenic proteins. In addition to 3D structural similarities, they calculate the sequence similarity between the input protein sequence and known allergenic proteins in the primary structure. The final allergenicity prediction is obtained by combining the results from these two complementary similarity-based approaches—3D structural and primary sequence alignments against known allergenic proteins [6].

#### 1.1.3 ProAll-D

This research has employed a biological feature extraction method called Amino Acid Composition (AAC), resulting in 125-dimensional feature vectors. We have incorporated this AAC-based feature extraction idea into our research experiments [7]. Specifically, we extracted AAC descriptors and combined them with various feature vectors to improve our prediction results. For classification, the authors of the ProAll-D have utilized Long Short-Term Memory (LSTM) networks— an architecture well-suited for sequence data such as proteins—to classify proteins as allergenic or non-allergenic [8].

#### 1.1.4 AlgPred 2.0

This research which has been referenced and benchmarked in many later works, has utilized biological and search-based approaches, combined with a machine-learning model, as a hybrid model. They extracted AAC descriptors and used them as feature vectors for various machine-learning models. The authors identified Random Forest [9] as the top-performing model among their evaluated machine-learning classifiers. Additionally, they published a dataset with individual training and validation sets, which dataset is used in our study, and the results on the validation dataset of the AlgPred 2.0 are used to compare recent work results [10].

#### 1.1.5 DeepAlgPro

This research has focused on the interpretability of the model. Since proteins are used in sensitive and important areas such as pharmaceuticals, the model must be reliable for pharmacologists and others who use proteins in new compositions. Ensuring the interpretability of the model can be highly valuable and useful, providing more reliability in the model’s predictions [11].

In this study, we have proposed a method called AllerTrans for recognizing allergenic and non-allergenic protein sequences. AllerTrans can classify these proteins with admissible accuracy, sensitivity, specificity, area under the ROC curve, and Matthews Correlation Coefficient (MCC). We combined various protein embedding vectors to improve model performance. We also effectively ensembled models together to achieve better results.

## 2. Methodology

In general, all the methods used in this research are comprised of the following two phases:

A. Feature Extraction
B. Training a Model on the Extracted Feature Vectors

In the first phase, we used Protein Language Models (pLM) to extract feature vectors from the primary structure of proteins presented in FASTA format. We trained classification models based on these feature vectors in the next phase. Our study involves binary classification, categorizing proteins as either allergenic or non-allergenic.

### 2.1 Feature extraction

This phase is crucial in laying the foundation for the following modeling phase. Two protein language models (pLM) were employed to extract embedding vectors from input protein sequences in FASTA format. These two models are ESM-2 [12] and ProtT5 [13]. One of the ideas for achieving better results is to combine extracted feature vectors. For this purpose, we utilized the concatenation of ProtT5 embedding vectors and ESM-2 embedding vectors. These concatenated embedding vectors proceed to the next phase and serve as the input for our classification model.

### 2.2 Modeling

A neural network designed for the binary classification task consists of three non-linear hidden layers with the Rectified Linear Unit (ReLU) activation function. Since this model is sufficiently simple, there is no need to employ techniques like dropout, weight decay, or other methods to prevent overfitting. According to the experiments, our best results are obtained by this simple neural network [14].

Various hyperparameter values were tested for this model, and after numerous experiments, the best result was achieved as described in Figure 1. The number of neurons in each hidden layer was a parameter that was determined through trial and error, and we found a combination that yielded the best result.

**Figure 1.**
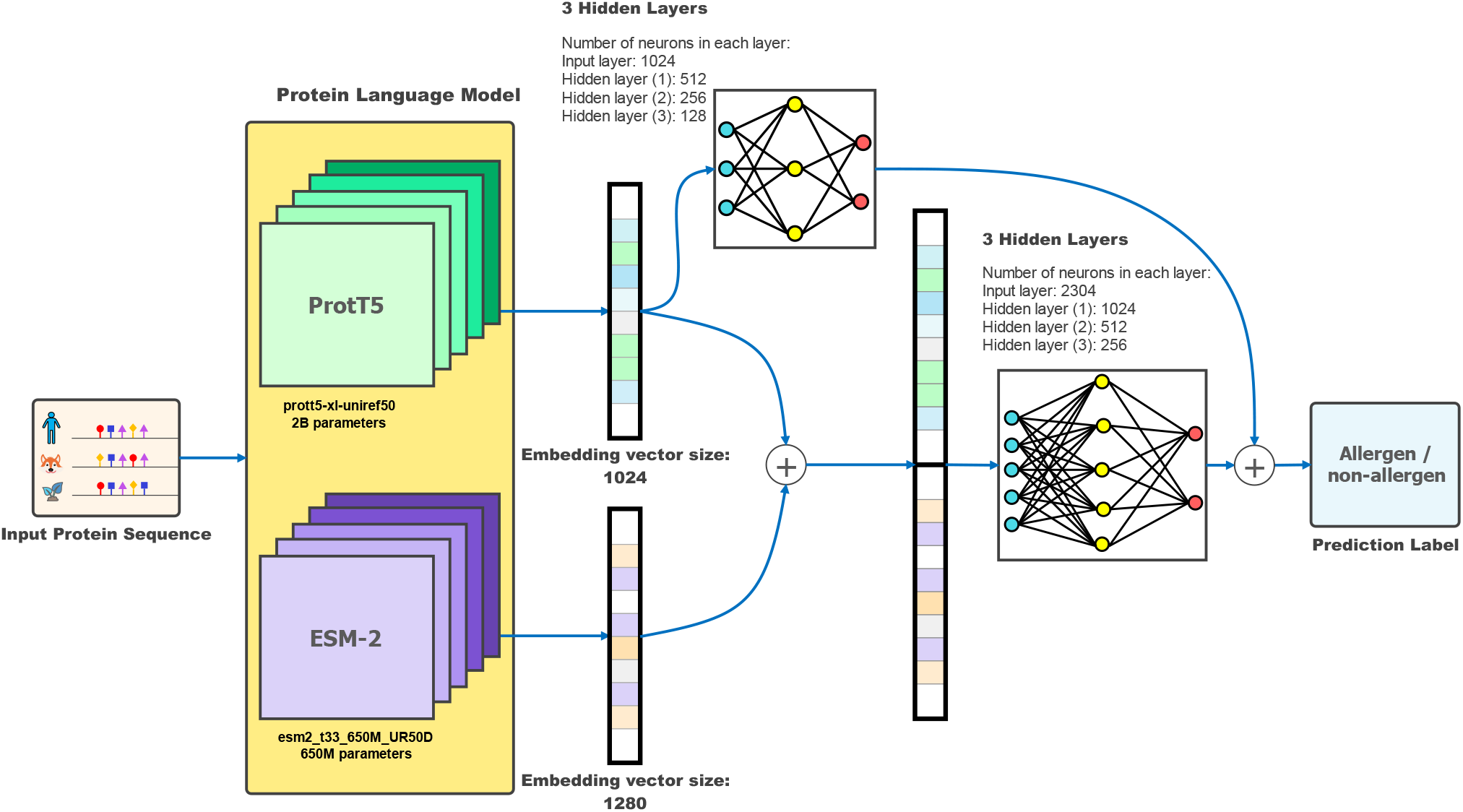
AllerTrans Architecture. The learning rate in our neural networks was 0.001 and they were trained in 180 epochs, with the loss function of Cross-Entropy, and optimizer of Adam.

### 2.3 Enhancement

Each type of feature vector represents a specific aspect of the protein sequence. Combining them through concatenation influences the final result based on their contributions. We noticed that the model exhibits higher specificity with ProtT5 embedding vectors alone. However, when the ProtT5 embedding vectors are concatenated with ESM-2, the sensitivity improves while the specificity decreases. Recognizing the importance of balancing sensitivity and specificity, we decided to effectively ensemble these two models. Using a single-layer neural network, we determined linear weights to combine the predictions of ProtT5 and ProtT-ESM, resulting in our best outcome.

### 2.4 Evaluation Metrics

We evaluate the models using the standard five-fold cross-validation method. In our study, we utilized the evaluation metrics outlined in the Raghava et. al. paper [10], for allergenic protein recognition models. As highlighted in that work, the most important evaluation metrics, along with the standard formulas that we used, are as follows:

#### 2.4.1 Sensitivity (Recall of the positive class)

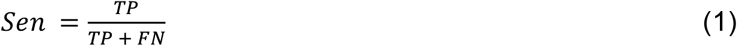

#### 2.4.2. Specificity (Recall of the negative class)

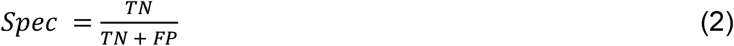

#### 2.4.3 AUC-ROC (Area under the ROC Curve) [15]

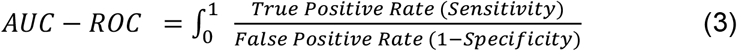

#### 2.4.4 MCC (Matthews Correlation Coefficient) [16]

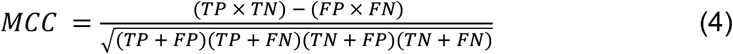

In the presented formulas, TP stands for the number of true positive predictions, TN stands for the number of true negative predictions, FP represents the number of sequences falsely predicted as positive, and FN indicates the number of sequences of false negative predictions.

## 3. Results & Discussion

Our final results are presented in Table 1, which provides a comprehensive overview of the performance of AllerTrans in comparison with other established methods for predicting protein allergenicity.

**Table 1.** Comparison of AllerTrans with existing methods.

| Method | Sens | Spec | Acc | AUC | MCC |
| --- | --- | --- | --- | --- | --- |
| **AllerTrans (5-fold)** | 97.91 | 97.69 | **97.80** | **0.99** | **0.956** |
| AllerTrans (AlgPred 2.0 test) | 93.5 | 95.38 | 94.44 | 0.98 | 0.9 |
| AlgPred 2.0 (test) | 89.3 | 95.09 | 92.26 | 0.98 | 0.85 |
| AlgPred | 88.87 | 81.86 | 85.02 | NA | 0.7053 |
| ALLERDET (AlgPred 2.0 test) | **98.46** | 94.37 | 97.26 | NA | NA |
| AllerCatPro 2.0 (AlgPred 2.0 test) | 93.2 | **98.8** | 96 | NA | 0.921 |
| AllerTOPv2 [17] | 86.7 | 90.7 | 88.7 | NA | 0.775 |
| AllerTOP [18] | 87.6 | 78 | 82.8 | NA | 0.671 |
| AllerHunter [19] | 83.7 | 96.4 | 95.3 | 0.928 + 0.004 | 0.738 |
| AllergenFP [20] | 86.8 | 89.1 | 87.9 | NA | 0.759 |

Several experiments were conducted, and ultimately, our most effective approach, AllerTrans, was chosen to predict the allergenicity of proteins.

In the feature extraction phase, we utilized two different approaches: biological methods and transformer-based language models [21] to extract feature vectors from protein sequences in FASTA format. Subsequently, classification models were trained based on these feature vectors. For the modeling phase, our classification models can be divided into two main categories: those based on classic machine-learning methods and those based on deep-learning methods. Figure 2 provides an overview of all the experiments conducted in our study.

**Figure 2.**
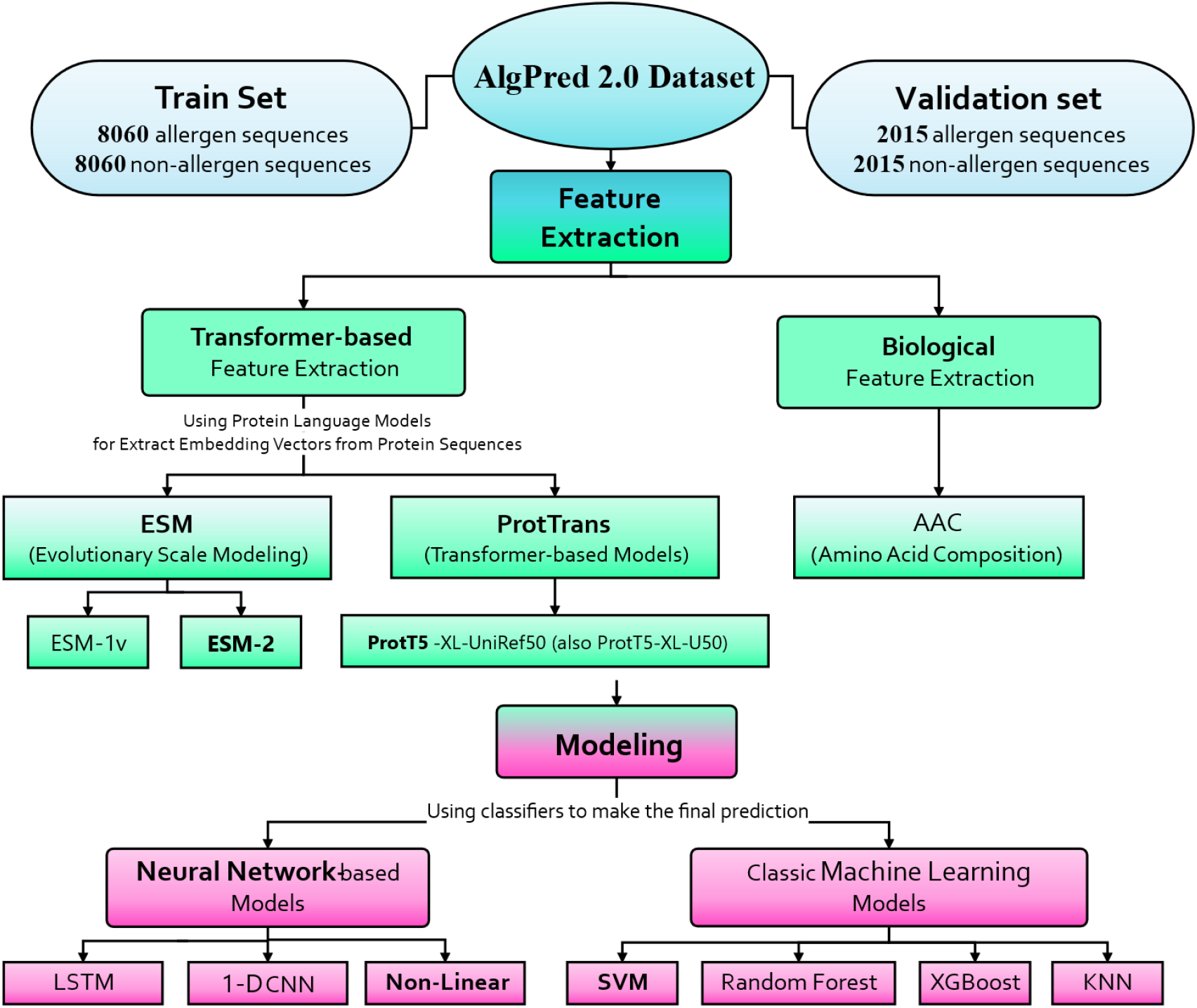
Overview of Experimental Approaches and their Relations in Protein Allergenicity Prediction.

We will discuss the different models, analyzing their performances and challenges. As an experiment, before classification with classic machine-learning models, we applied an autoencoder [22] to the embedding vectors and then conducted modeling for binary classification. For deep learning-based methods, the dimensionality reduction of the input can be replaced by adding layers in the neural network, instead of employing a separate autoencoder.

After the initial modeling, to achieve better results, we utilized hyperparameter tuning methods to build the best possible model. Finally, since it’s anticipated that appropriate leveraging of ensemble models can yield improved outcomes, we merged the predictions using two ensemble methods.

The idea of the first method was to combine the results of two models by considering their confidence levels. When selecting the result, if both models agreed, the outcome was determined. However, if we wanted to combine the results of two models, while one predicted class 1 (allergen) and the other predicted class 0 (non-allergen), we examined the model outputs before the thresholding stage and selected the final result based on the model that had more confidence in its prediction. The second method for ensemble models required determining the weights of different models. The impact of distinct models could vary, and to find the optimal weights for the models, we utilized a simple single-layer neural network. The final result was determined based on the second approach for ensemble models.

As highlighted in the introduction section, the models developed for allergenicity prediction play a critical role in assessing the safety of proteins, particularly recombinant proteins. Incorrectly categorizing an allergenic protein as non-allergen can pose serious risks in medical drug development. For instance, a medical drug containing a recombinant protein, incorrectly labeled as non-allergen while being allergenic, has the potential to cause harm to allergic patients, and jeopardize their health due to the misidentification of the protein’s allergenicity. Therefore, the Sensitivity metric is one of the most crucial in this area of research and serves as one of the most important metrics for selecting our final model.

### 3.1 Experimental Results

We started by employing the ESM-2 model to extract embedding vectors, and for classification, we initially used classic machine-learning models. The results of these classifiers are shown in Table 2. All the subsequent results are derived from training on the AlgPred 2.0 train set and evaluating the performance on the AlgPred 2.0 validation set.

**Table 2.**
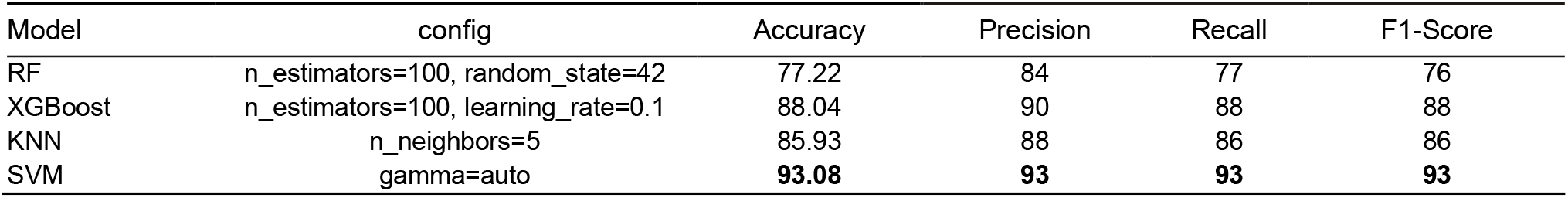
Comparison of classic machine-learning classifiers on ESM-2 embedding vectors.

These initial results demonstrate that the Support Vector Machine (SVM) model achieves better outcomes compared to other classic machine-learning models including Random Forest (RF), eXtreme Gradient Boosting (XGBoost) [23], and K-nearest neighbor (KNN) [24]. The reason is highly attributed to its high generalization capability and superior ability to handle high-dimensional data. Given the substantial number of features—here, 1280 is the embedding vector size of our leveraged ESM-2 model—the SVM model outperforms other classical machine-learning models [25]. Subsequently, we proceeded with tuning the hyperparameters of the SVM model and other models, and the results after finding the optimal hyperparameters are presented in Table 3 [26].

**Table 3.** Comparison of classic machine-learning classifiers with optimal hyper-parameters on ESM-2 embedding vectors.

| Model | config | Accuracy | Precision | Recall | F1-Score |
| --- | --- | --- | --- | --- | --- |
| XGBoost | n_estimators=100, learning_rate=0.001, gamma=0.1 | 88.78 | 89 | 89 | 89 |
| KNN | n_neighbors=10 | 86.75 | 89 | 89 | 89 |
| SVM | kernel=rbf, gamma=0.01, C=0.5 | **94.07** | **94** | **94** | **94** |

Subsequently, we utilized an autoencoder to reduce the input dimensionality of the model. Once again, classic machine-learning models were applied for classification. The impacts of employing an autoencoder before classification are detailed in Table 4.

**Table 4.** Comparison of classic machine-learning classifiers on dimensionality-reduced ESM-2 embedding vectors with Autoencoder.

| Model | config | Accuracy | Precision | Recall | F1-Score |
| --- | --- | --- | --- | --- | --- |
| RF | n_estimators=100, random_state=42 | 88.31 | 90 | 88 | 88 |
| XGBoost | n_estimators=100, learning_rate=0.1 | **92** | **91** | **91** | **91** |
| KNN | n_neighbors=10 | 87.87 | 89 | 89 | 89 |
| SVM | gamma=auto | 50.57 | 75 | 51 | 35 |

The autoencoder positively impacted the performance of all experimented machine-learning models, except for SVM. As anticipated, the SVM model, known for its commendable generalization in handling high-dimensional inputs experienced a significant decrease in performance. The reduction in input dimensions led to missing some information, posing a challenge for the SVM model to perform effectively. Additionally, we employed single-layer Long Short-Term Memory (LSTM) [27] and 1-dimensional Convolutional Neural Network (1D-CNN) [28] as classifiers, and the results are presented in Table 5.

**Table 5.** Comparison of classifier models on ESM-2 embedding vectors.

| Model | Accuracy | Precision | Recall | F1-Score |
| --- | --- | --- | --- | --- |
| Single-layer LSTM | 93.33 | 93.46 | 93.33 | 93.32 |
| 1D-CNN | 84.69 | 84.31 | 84.26 | 82.22 |
| SVM | **94.07** | 94 | 94 | 94 |
| RF + Autoencoder | 88.31 | 90 | 88 | 88 |
| XGBoost + Autoencoder | 92 | 91 | 91 | 91 |
| KNN + Autoencoder | 87.87 | 89 | 89 | 89 |
| Non-Linear NN | 94.04 | **94.07** | **94.04** | **94.04** |

The last row of this table represents a multi-layer perceptron with the following configuration:
- Loss Function: Cross Entropy Loss - Optimizer: Adam - Learning Rate: 0.001 - Number of Epochs: 150 - Input Layer: 1280D - Hidden Layer 1: 200D - Hidden Layer 2: 100D - Hidden Layer 3: 50D

We developed neural network-based models using PyTorch [29] and classic machine-learning models using scikit-learn [30].

After this stage, we employed a non-linear neural network with the architecture introduced in the previous section as the classifier in all subsequent experiments. We also extracted embedding vectors using the ProtT5 model. In another experiment, we employed the biological rule-based method for feature extraction named Amino-Acid Composition (AAC). This method of feature extraction singularly did not improve outcomes. However, concatenating the feature vectors extracted by AAC with the embedding vectors obtained from the ProtT5 model led to improvement. This is because the biological method can extract some solid features, and concatenating with ProtT5 can introduce more features and information, resulting in good performance for binary classification. Given the numerous experiments, we will showcase only the best results of each technique. Subsequent experiments involved modifying and concatenating feature vectors, employing ensemble methods, and hyperparameter tuning.

We utilized the five-fold cross-validation method for evaluating our models, and the comparison between our top-performing models and Algpred 2.0 results of five-fold cross-validation are reported in Table 6. The model naming convention ‘X’ + DNN indicates that we obtained feature vectors from ‘X’ and then input them into our defined non-linear Deep Neural Network (DNN). In this context, ‘X’ can refer to either a single feature extraction method or a combination of different feature extraction methods.

**Table 6.** Performance evaluation of top models through five-fold cross-validation & comparison with Algpred 2.0 results.

| Method | Sens | Spec | Acc | AUC | MCC |
| --- | --- | --- | --- | --- | --- |
| Algpred 2.0 (five-fold) | 93.10 | 95.36 | 92.26 | 0.99 | 0.88 |
| ProtT5 + DNN | 92.1 | **96.53** | 94.3 | **0.99** | 0.8887 |
| Prot-ESM + DNN | **95.86** | 93.28 | **94.59** | 0.9839 | **0.8925** |

The results presented in Table 6 have motivated us to effectively ensemble two models—Prot-ESM and ProtT5—to improve both the sensitivity and the specificity, achieving a better balance between them. Table 7 compares our top-performing models and the AlgPred 2.0 model evaluation on the AlgPred 2.0 test set.

**Table 7.**
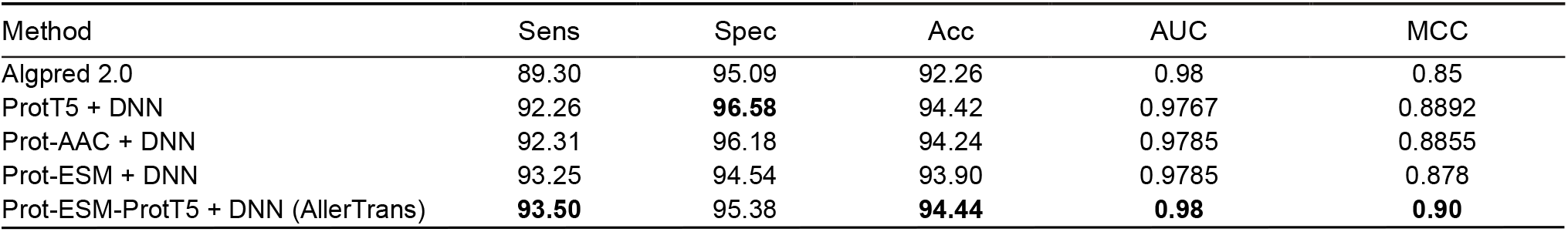
The final comparison of our top models on the AlgPred 2.0 validation set.

| Method | Sens | Spec | Acc | AUC | MCC |
| --- | --- | --- | --- | --- | --- |
| Algpred 2.0 | 89.30 | 95.09 | 92.26 | 0.98 | 0.85 |
| ProtT5 + DNN | 92.26 | **96.58** | 94.42 | 0.9767 | 0.8892 |
| Prot-AAC + DNN | 92.31 | 96.18 | 94.24 | 0.9785 | 0.8855 |
| Prot-ESM + DNN | 93.25 | 94.54 | 93.90 | 0.9785 | 0.878 |
| Prot-ESM-ProtT5 + DNN (AllerTrans) | **93.50** | 95.38 | **94.44** | **0.98** | **0.90** |

### 3.2 Comparative Analysis of Results with Related Work

As reported in Table 1, the current state-of-the-art models in protein allergenicity prediction are AllerCatPro 2.0 and AllerDET. Our results demonstrate an improvement over these two models, emphasizing the importance of maintaining a balance between sensitivity and specificity metrics for the protein allergenicity prediction model. This balance is also reflected in the AUC-ROC metric, which is 99% for AllerTrans using five-fold cross-validation on the AlgPred 2.0 dataset. Notably, the specificity of AllerTrans surpasses the specificity of AllerDET, and the sensitivity of AllerTrans outperforms AllerCatPro 2.0.

Moreover, as the approach of AllerCatPro 2.0 is described in the introduction section, we notice that although this research is precious, it is very complex and requires space for saving allergenic protein sequences and also the 3D structures of well-known allergens, or searching for them online, which is time-consuming when searching for many allergen proteins. Additionally, the model’s performance is intricately linked to these well-known allergenic proteins, demanding periodic updates as new allergens are identified.

Furthermore, upon reviewing the approach outlined in the introduction section for AllerDET, we recognize the value of this research. However, the similarity-based approach in this study is highly time-consuming and has limitations of similarity-based approaches as we described them in the introduction section. To extract the demonstrated four features for the AllerDET model, they find similarities between the query protein sequence and all protein sequences in the train set, which requires considerable time.

### 3.3 Data-Driven Analysis (DDA)

A data-driven analysis was conducted in our research to determine the major reasons for incorrect predictions. We utilized the BLAST method and the UniProt database [31] to investigate the properties of the incorrectly labeled sequences. The results were intriguing. The incorrectly labeled sequences mostly had an identity of less than 100% in UniProt, indicating that the exact protein sequence isn’t available on the UniProt website. Our utilized protein language models—ESM and ProtT5—were trained on the UniProt dataset. Therefore, they may have limitations in accurately predicting the embedding vectors of protein sequences not identified in the UniProt database.

### 3.4 Feature Work

Our next goal could be to use knowledge distillation to train a network named RWKV [32] as the student network, with a transformer-based protein language model as the teacher network. This idea can be valuable because Recurrent Neural Networks (RNN) may have better prediction time than transformer-based models. Therefore, it would be invaluable to train a network using this technique to achieve performance similar to that of a transformer-based network, but with faster prediction times.

## 4. Conclusions

In this research study, we aimed to develop the best possible deep-learning model for predicting protein allergenicity. Throughout the study, we experimented with various methods, compared the results at each stage, and proposed ideas for improving the model, based on the obtained outcomes. Finally, the best results were achieved with the “Prot-ESM-ProtT5 + DNN” approach. Our final model demonstrated improvements across all evaluation metrics compared to AlgPred 2.0. The results of the evaluation metrics have improved by up to 5%. Achieving improvements in high values of evaluation metrics—here the previous research studies demonstrated higher than 90 values in most evaluation metrics—is challenging, making even a slight enhancement valuable. Although we had other top models, we selected the “Prot-ESM + ProtT5” model for AllerTrans because, in this context, the sensitivity metric and the balance between sensitivity and specificity take priority.

## Data Availability

The source code and evaluation results are available at the AllerTrans public repository: https://doi.org/10.6084/m9.figshare.26524561.v1.

This research study utilized the publicly available AlgPred 2.0 train and validation sets: https://webs.iiitd.edu.in/raghava/algpred2/stand.html.

